# Gene expression adaptation of metastases to their host tissue

**DOI:** 10.1101/2024.12.18.629130

**Authors:** Luise Nagel, Marten Wenzel, Sascha Hoppe, Patrick S. Plum, Mohammad Karimpour, Marek Franitza, Roger Wahba, Marc Bludau, Christiane J. Bruns, Alexander Quaas, Andreas Beyer, Axel M. Hillmer

## Abstract

The adaptation of metastatic cells to their host tissue critically determines the pathogenicity of a cancer and therefore patient survival. Yet, it remains elusive to what extent the host environment drives gene expression programs in metastatic cells. Here we identify adaptive mechanisms that enable metastases to establish themselves in a novel tissue context. We performed single-cell RNA-sequencing on malignant and benign tissue samples from untreated donors with colorectal adenocarcinoma and liver metastasis to deduce tissue adaptive expression patterns. A novel computational approach identified genes and pathways that consistently adapted to the host tissue at the transition from the primary tumor to the paired metastasis across donors. This analysis revealed that the majority of expression changes in the metastasis reflect an expression signature reminiscent of benign liver epithelial cells. Cellular processes adapting to the liver environment include basic cellular functions such as energy metabolism, as well as tissue-specific pathways such as the regulation of lipid metabolism by PPAR-α. These adaptations potentially increase the pathogenicity of the metastatic cells and may provide new therapeutic strategies.

## Introduction

The formation of distant metastases necessitates cancer cells to leave the primary cancer site, travel as a circulating tumor cell to another organ, infiltrate this organ and adapt to the novel surrounding tissue and grow as a metastasis^1–3^. These obstacles result in less than 0.01% of the primary tumor cells metastasizing^4^. The fact that a significant number of cancer cells can circulate in the bloodstream but forming a limited number of metastases, known as the ineffectiveness of metastatic spread^5–8^, illustrates the barrier cancer cells have to overcome to survive and proliferate in a novel tissue context. Given that metastases are responsible for over 90% of cancer-related deaths, enhancing our understanding of this process is of great importance to increase patient survival^9^. In the 19th century, Stephen Paget suggested the ‘seed and soil’ theory saying that a cancer cell (seed) requires a specific tissue (soil) to form a metastasis^10^. Further, it was recognized that patterns of metastases are influenced by architecture of tissues and blood stream^11^. While both factors are valid and contribute to the likelihood that a metastasis forms, the ability of a cancer cell to adapt is an additional pivotal factor for the establishment of metastases.

The specific gene expression signature of metastatic cells is determined by four factors: (1) the tissue of origin, (2) the cancer-specific expression signatures^12–16^, (3) the metastatic process and (4) adaptation to the host tissue. Expression patterns of metastases show a strong similarity to that of the primary tumor they originate from^17^, through the presence of two key characteristics: (1) resemblance to the expression signature of the original cell type from which the tumor arose and (2) cancer-specific alterations. Metastasis-specific processes such as mesenchymal-epithelial transition (MET) and epithelial–mesenchymal transition (EMT), including dissemination, travel, invasion^1,13,18–21^, as well as escaping the immune system^22–24^, necessitate the involvement of specific pathways and functions, resulting in an (3) altered gene expression pattern that is distinct from that of the primary tumor^1,18^. Furthermore, the expression profile of the metastases is driven by (4) the novel surrounding tissue in which they establish themselves, as cancer cells undergo a tissue-specific metabolic adaptation process to integrate and survive in a new organ^25^. Metastases need to display a high metabolic flexibility, enabling them to accommodate for different environments with different oxygen abundances, energy sources, and nutrient availability^26,27^. Importantly, while this adaptation may be enhanced or ‘helped’ by genetic and/or epigenetic predisposition in metastasising cells, such (cell-intrinsic) predisposition is not sufficient to explain all metastasis-specific processes. Instead, this adaptation also results from cell-extrinsic signals in the new environment^28^, such as local nutrient availability and paracrine signaling resulting from tissue-resident cells. In addition to explaining the establishment of metastases in their new environment, this adaptation may also modulate the pathogenicity of metastases independent of their genotype. Thus, better understanding the interaction between metastases and their environment is of utmost clinical importance.

Monitoring gene expression in metastatic cells has a great potential for elucidating the adaptive processes enabling metastasising cells to successfully establish themselves in the new tissue environment. However, earlier work has mostly focussed on the direct comparison of gene expression differences between metastases and their primary tumors without addressing the question to what extent those differences may be driven by metabolic adaptation^13,18,19,21^. Other research has focussed on the direct interaction between the metastasis and its microenvironment with a specific focus on immune cells^22–24^. Although this research is highly relevant for understanding immune evasion, it does not address the question of metabolic tissue adaptation.

Here, we present a new combined experimental and bioinformatic strategy for the identification of tissue-adaptive genes and pathways, i.e. genes/pathways whose altered expression in metastases reflects an adaptation to the new host tissue. Our strategy is based on the simultaneous single-cell profiling of four cellular compartments from the same donor: primary tumor cells, tumor cells of the metastasis and benign cells of the two respective host organs. Importantly, benign cells are sampled distant from the tumor sites to minimize direct influence of the tumor on them. Using a bioinformatic approach we identify expression differences between the metastasis and the primary tumor matching those observed between the two healthy tissues.

Once cancer has developed metastases and turned into systemic disease, surgery with curative intent is usually not beneficial anymore resulting in a scarcity of clinical specimen suitable for single cell sequencing approaches. Colorectal cancer (CRC) that formed a limited number of distinct detectable synchronous metastases in the liver belongs to the rare clinical situations where, despite metastases, surgery increases chances for prolonged survival and therefore is used in the clinic^29,30^. Usually, however, neoadjuvant or perioperative therapy is applied before surgery to such patients in Germany which might lead to persisting cell biological processes in the metastases. We have deliberately selected patients without neoadjuvant treatment and simultaneous surgery on colon and liver to get an unbiased view on the cell states and respective transcriptomes of colorectal cancer cells of the primary tumor and metastases within the same patient together with cells of the surrounding normal tissues.

Using primary colon cancer with liver metastases as a model system, we analyze the adaptation of cancer cells to their novel environment. This analysis reveals that the majority of expression changes in the metastasis reflect an expression signature reminiscent of benign liver epithelial cells. Cellular processes adapting to the liver environment include basic cellular functions such as energy metabolism, as well as tissue-specific pathways such as the regulation of lipid metabolism by PPAR-α. These adaptations potentially increase the pathogenicity of the metastatic cells and may provide new therapeutic strategies based on inhibiting the metabolic adaptation of cancer cells to their host environment.

## Results

### Single-cell RNA sequencing of paired colon cancers and liver metastases

Matching tissue from five donors with untreated CRC and liver metastasis were sampled and underwent scRNA-seq (Figure 1a, b, Extended Data Table 1, see Methods). We performed scRNA-seq on tumor samples (which consist of a mix of benign and tumor cells) and of healthy tissue distal from the tumor. After stringent quality control, a total of 18,312 cells of 16 tissue samples were obtained (Figure 1c, Extended Data Table 2), which after normalization and integration were clustered into 30 clusters (Extended Data Figure 1d, Extended Data Table 2). Differential expression analysis was applied to identify cluster-specific marker genes (see Methods) and cell types were annotated utilizing previously known cell type-specific expression patterns (Figure 1d). Clusters commonly consisted of cells from different donors (Extended Data Figure 1a, b, c) and 20 of 23 clusters were composed of cells from liver and colon (Figure 1d). The most frequent cell type found across all tissues were fibroblasts, making up the majority of cells in both the healthy and tumor colon samples.

**Figure 1:**
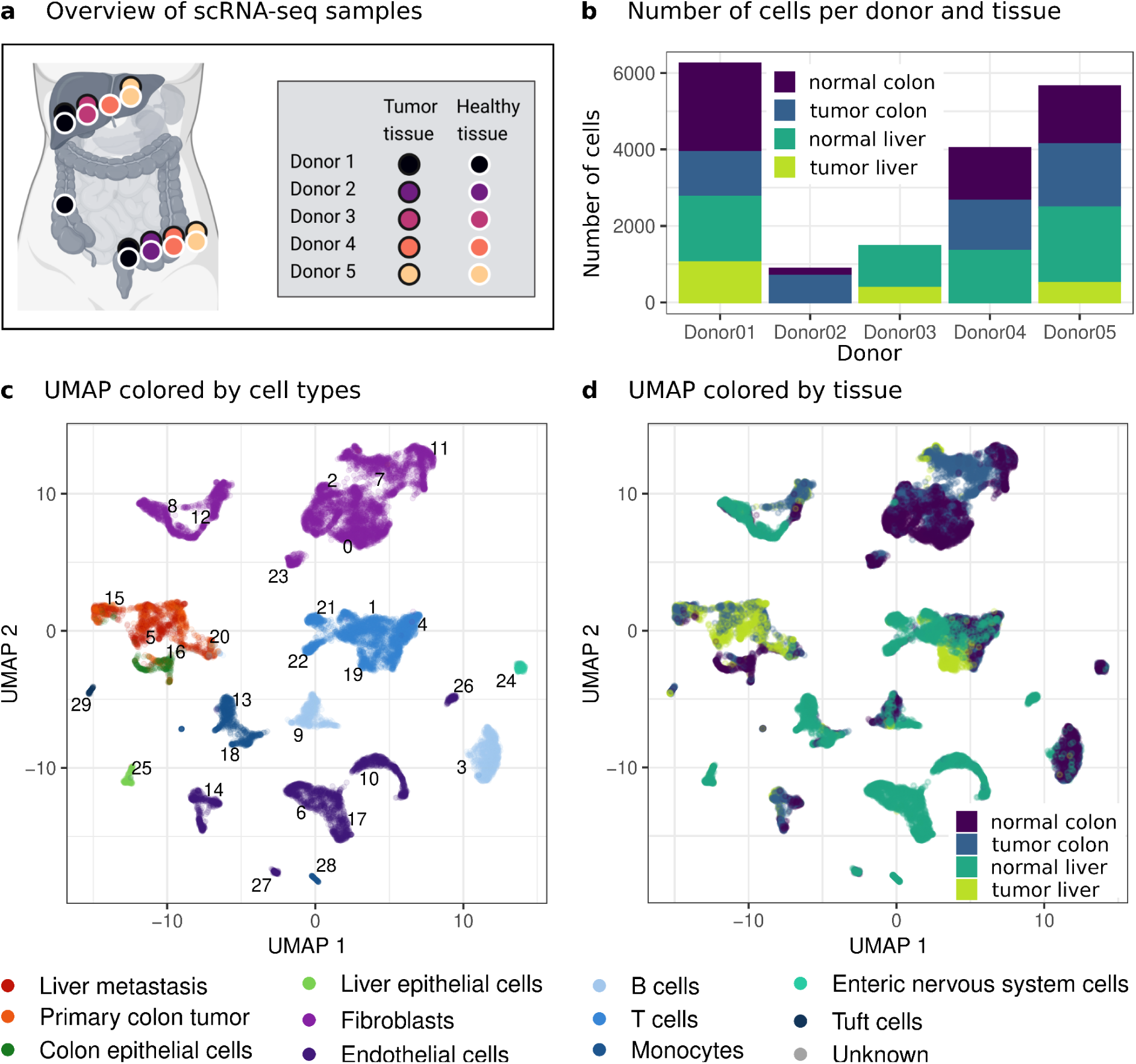
Overview of our scRNA-seq dataset from five donors with primary colon tumor and liver metastasis. **a)** Overview of scRNA-seq samples available from each donor (Partly created with biorender.com.) **b)** Number of cells per donor colored by sample origin site (normal colon (purple), tumor colon (blue), normal liver (turquoise), tumor liver (light green). **c)** UMAP of scRNA-seq data with colors indicating different cell types. Cluster numbers are shown in the plot. **d)** UMAP of scRNA-seq data with colors indicating different sample site origins. Colors as in b).

### Differentially expressed genes between primary tumor and metastasis cells

For the identification of differences in the gene expression between cancer cells from the primary tumor versus cancer cells from the metastasis, we searched for genes that are consistently higher in the primary tumor or metastasis across multiple donors. Using all tumor cells from donors where both primary tumor and metastasis was sequenced, we estimated donor-specific pseudo-bulk expression profiles and fold changes for each sample. Genes that were overexpressed in either the metastasis or the primary tumor across all donors were defined as ‘metastasis specific’ (MT-specific; 305/6872) or ‘primary tumor specific’ (PT-specific; 530/6872 genes), respectively (Figure 2a, 2b, Extended Data Table 3, see Methods). A small number of genes (131/24642) were found to be expressed in a tissue-exclusive manner, as they were virtually undetectable in the respective other tissue (primary tumor exclusive genes: 49/24642, metastasis exclusive genes: 82/24642, Extended Data Table 5, see Methods).

**Figure 2:**
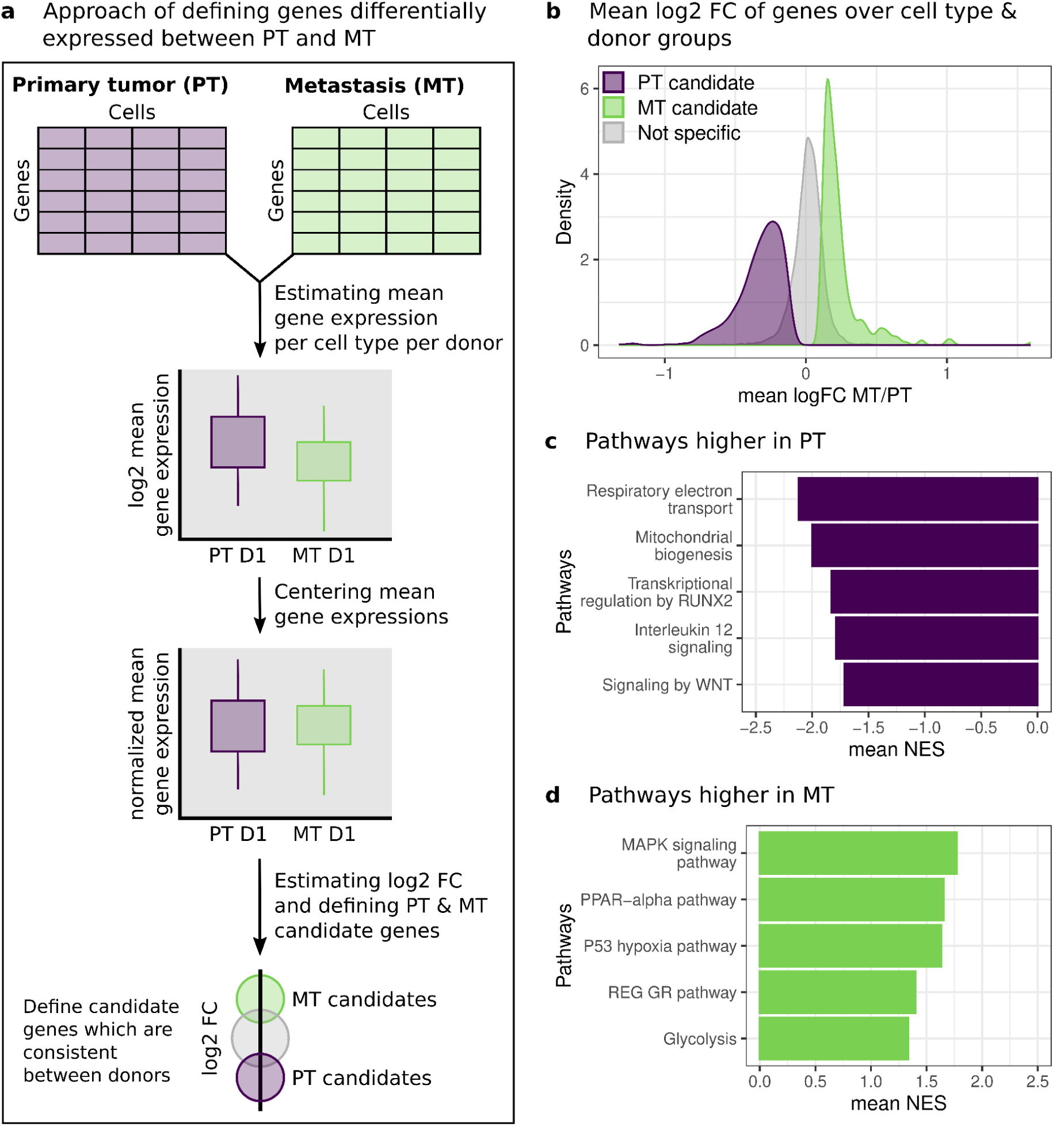
Gene expression differences between metastasis and primary tumor cells. **a)** Computational approach of defining differentially expressed genes between MT and PT in different donors. Exemplified here for donor 1 (D1). **b)** Average log2 fold change (FC) of centered mean gene expression over donors between MT and PT. PT specific (purple), MT specific (green) and non-differentially (gray) expressed genes are plotted individually. **c)** Five of the differentially expressed pathways in PT cells when compared to MT cells. The mean normalized enrichment score (NES) across the donors is displayed. **d)** Five of the most upregulated pathways in MT cells compared to PT cells. Mean NES across the donors is displayed.

Many of the genes found to be differentially expressed between the primary tumor and metastasis can be linked to the metastatic process, such as the cysteine-rich intestinal protein (*CRIP*), which is upregulated in the metastasis cells. *CRIP1* is a known oncogene, and has been observed to be higher expressed in tumors than in the tumor-adjacent tissues^31^. *CRIP1* also plays a role in cell migration and invasion in various cancer types including colorectal cancer. *CRIP1* mediates the epithelial-mesenchymal transition (EMT) and is therefore directly involved in the metastasis formatting processes.^32,33^ Our analysis also revealed some new candidate genes that are potentially involved in metastasis formation. For example, Fatty Acid-Binding Protein 1 (*FABP1*) was consistently higher expressed in the liver metastasis compared to the primary tumor. *FABP1* is mainly expressed in the liver and intestine, and as an intracellular protein involved in the binding, transport and metabolism of long-chain fatty acids^34,35^. Elevated *FABP1* expression has been reported for various tumor types, including colorectal and hepatocellular carcinomas^36^. Although *FAPB1* has not yet been directly associated with metastasis formation, exogenous *FABP* expression has, however, been associated with tumor progression and invasiveness^37^.

Pathway enrichment analysis was additionally performed, providing an overview of the cellular processes differentiating between the primary tumor and metastasis (Figure 2, Extended Data Table 5). This analysis revealed differences in the energy metabolism between the primary and secondary tumor. While glycolysis was upregulated in the metastasis, genes involved in oxidative phosphorylation (respiratory electron chain & mitochondrial biogenesis) were more highly expressed in the primary tumor. This is consistent with previous reports, showing that glycolytic dominant metabolic reprogramming commonly occurs during the metastatic process, resulting in an increase in glycolysis in the metastasis compared to the primary tumor^26,38,39^. The switch from oxidative phosphorylation towards glycolysis has been described as the Warburg effect^40^, a metabolic phenotype characteristic for highly proliferative cancer cells. Cells adapt their energy metabolism to the environment. For example, hypoxia is known to drive cells into glycolysis and consistent with that notion we observed a slight upregulation of Hypoxia Inducible Factor 1 Subunit Alpha (*HIF1A*) in metastatic cells.^41,42^

### Adaptation of gene expression to the novel tissue context

Next, we aimed to test if metastatic cells may adapt their metabolism to the new liver environment. Note that the peroxisome proliferator-activated receptor alpha (PPAR*αα*) pathway was upregulated in the metastatic cells compared to the primary tumor (Figure 2d). This observation could serve as a first hint towards a metabolic adaptation of metastatic cells to the liver environment, since PPAR*αα* signaling plays a key role in hepatocyte biology^43,44^. We expected that the expression patterns of ‘tissue-adaptive genes’ would agree between metastatic colon cancer and normal liver cells as both reside in the same environment. To test this hypothesis we focused on epithelial cells, because the primary tumors arose from colon epithelial cells and epithelial cells are the main cell type present in the liver, where metastasis manifested. When comparing gene expression in tumor cells to their surrounding epithelial cells we noted that the majority of PT-specific genes were higher expressed in healthy colon compared to healthy liver, while MT-specific genes were more highly expressed in healthy liver compared to healthy colon (Figure 3a). Thus, the majority of differences in the gene expression between the primary and secondary tumors are reflected in the surrounding tissue, strengthening the hypothesis that the surrounding tissues may drive expression patterns in the tumors.

**Figure 3:**
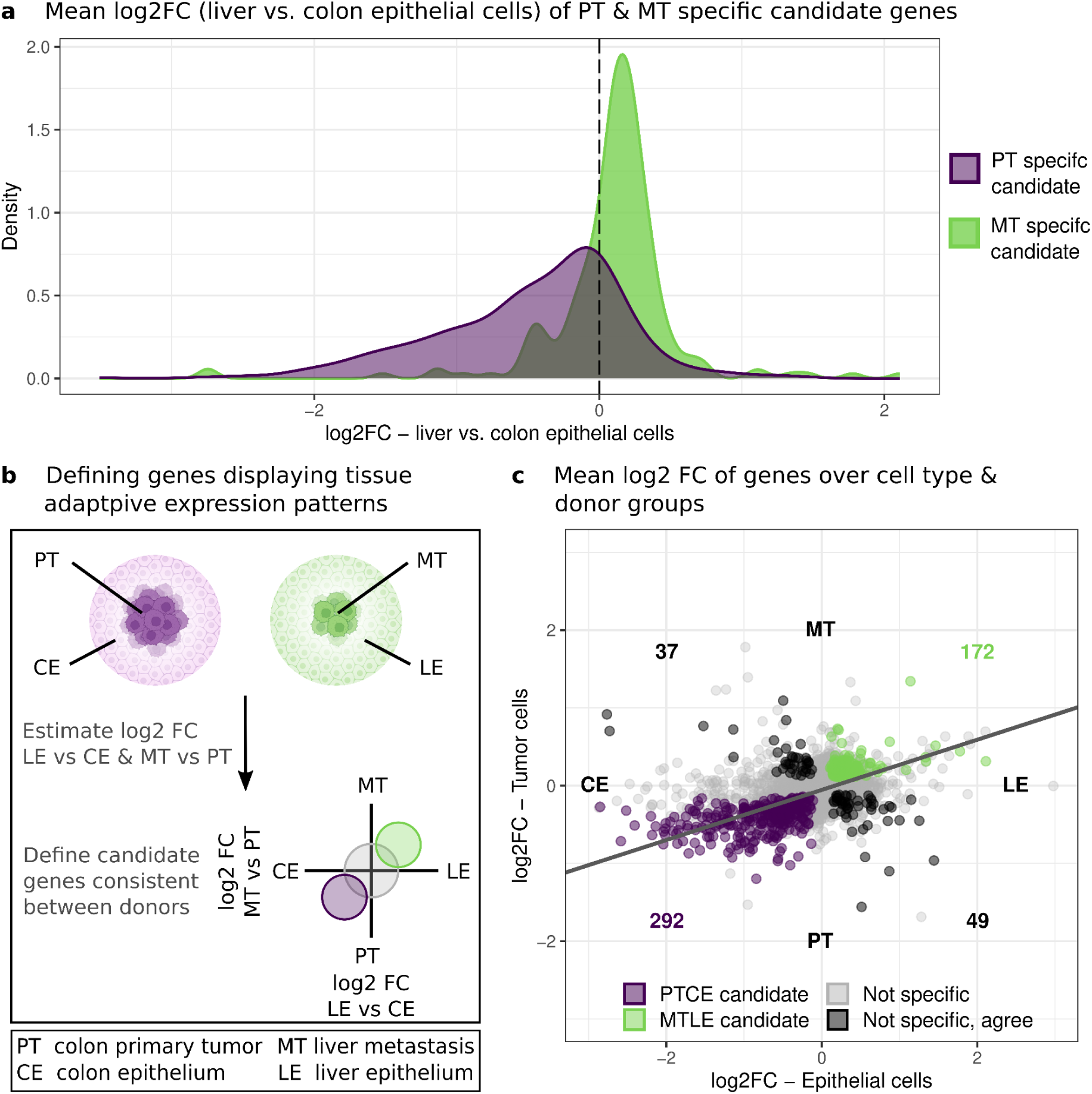
Gene expression differences between metastasis and primary tumor as well as liver and colon epithelial cells. **a)** Mean log2 fold change of liver to colon epithelial cells of genes defined as primary tumor (purple) or metastasis (green) specific in 3a and b. **b)** Bioinformatic pipeline of defining genes which are differentially expressed between metastasis and primary tumor as well as liver and colon epithelial cells in different donors. **c)** Mean log2 fold change of metastasis and primary tumor plotted against the mean log2 fold change of liver and colon epithelial cells estimated over the donors. Genes with expression patterns consistent between the donors are highlighted (primary tumor and colon epithelial specific genes = purple, metastasis and liver epithelial specific genes = green, other = black, four outlier genes are not visualized in the plot). The deming regression coefficient is displayed as a gray line.

To systematically identify tissue-adaptive genes, we compared expression differences between the metastasis and the primary tumor to expression differences between normal liver and normal colon epithelial cells (Figure 3b). We termed a gene ‘adaptive’ either if its expression was upregulated in the metastasis (MT) relative to the primary tumor (PT) *and* more highly expressed in the liver epithelial (LE) compared to the colon epithelial (CE) cells, or alternatively, if the gene was downregulated in the metastasis and accordingly more lowly expressed in healthy liver compared to healthy colon. Thus, we computed respective expression fold changes per donor (to control for donor-specific expression patterns), and classified genes into primary tumor & colon epithelial (PTCE) or metastasis & liver epithelial (MTLE) specific based on consistent fold changes across the donors used within the analysis (Figure 3b, Extended Data Table 4, see Methods). Some genes were tissue-exclusive, i.e. undetectable in some of the tissues, but expressed at relevant levels in others (Extended Data Table 5, see Methods).

Out of the 5,019 genes analyzed, 550 (i.e. ∼11%) had consistent expression changes across donors (Figure 3c, Extended Data Figure 2b, c). Of those, the vast majority was either consistently upregulated in the primary tumor and colon epithelial cells (292 PTCE genes), or in the metastasis and liver epithelial cells (172 MTLE genes, Figure 3c). Thus, these 464 genes (84% of 550) had expression changes in the metastasis qualitatively matching expression differences in the respective healthy host tissues. Only a small fraction of genes (n = 37, n = 49) consistently (i.e. across donors) changed their expression in opposing directions (49 genes upregulated in primary tumor and liver epithelial, 37 genes upregulated in metastasis and colon epithelial). This result reveals an alignment between the expression patterns of tumor cells and their surrounding tissue. We note that the PTCE and MTLE genes do not contain all PT and MT specific genes defined above (Figure 2b) due to small differences in which genes are included in the analysis (see Methods). These changes in the preprocessing however only result in minimal differences in the analysis and do not change the overall results (Extended Data Figure 3a, b, c).

We also noted that the expression fold changes in the tumor cells (MT versus PT) were on average smaller than the fold changes in the healthy tissues (LE versus CE; Figure 3c). This finding is suggesting that metastatic cells move towards a ‘liver-like’ state, but do not fully adopt the expression program of liver epithelial cells.

### Validation of tissue adaptive genes in external data sets

To validate the PTCE and MTLE genes, we utilized an external scRNA-seq with primary colon tumor and matching liver metastasis cells.^45^ Pseudo-bulk log fold changes were estimated between the metastatic and primary tumor cells (Methods; Figure 4a, 4b, Extended Data Figure 4a). The MTLE genes commonly displayed a positive log fold change in the tumor and epithelial cells, whilst the majority of the PTCE genes had a negative log2 fold change in the external scRNA-seq data (328 out of 422 genes measured in both data having the same direction of effect).

**Figure 4:**
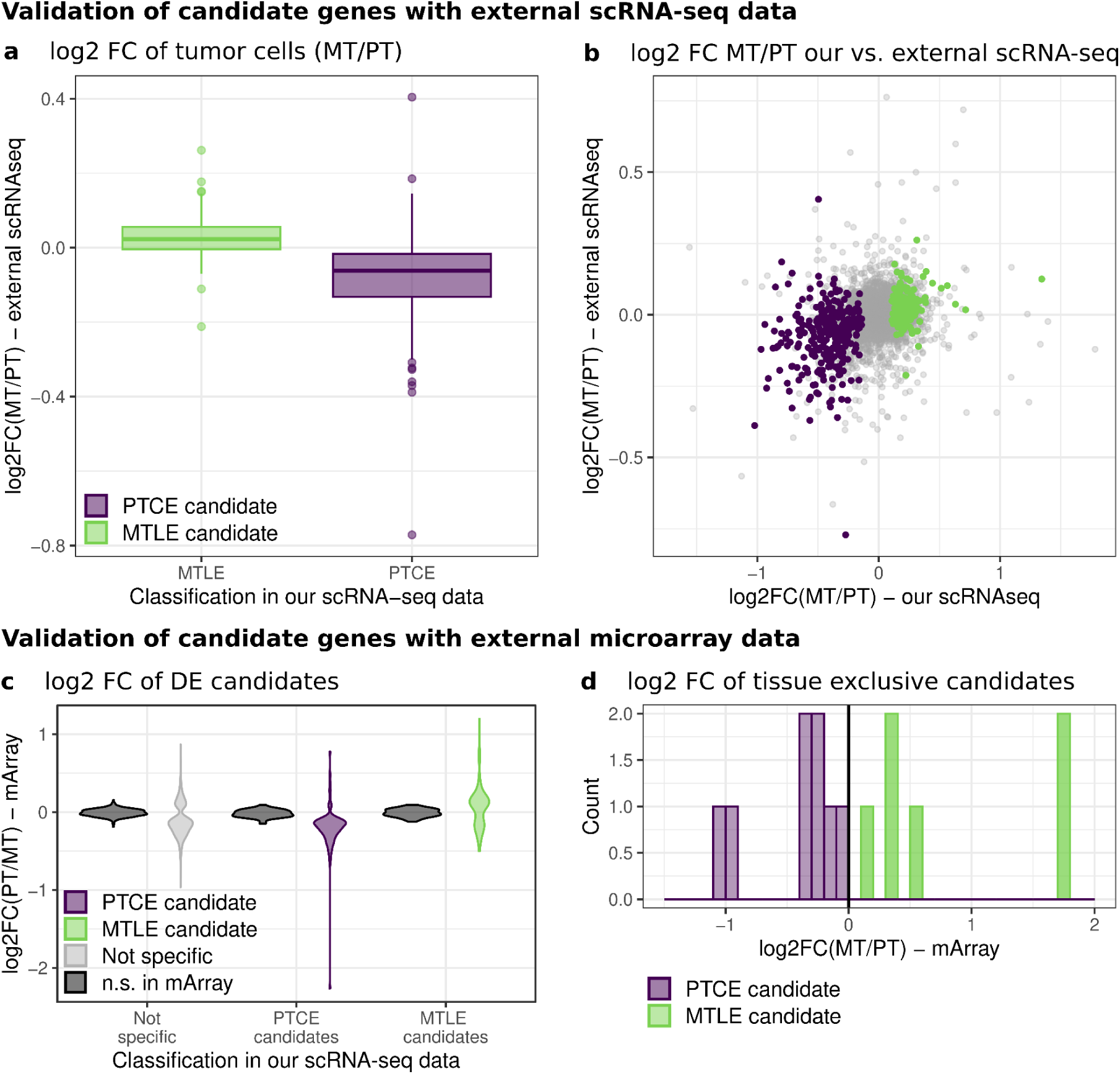
Validation of MTLE and PTCE candidate genes with external data. **a)** Log2 FC from metastasis to primary tumor cells in external scRNA-seq data plotted by their classification in our data (PTCE candidates = purple, MTLE candidates = green). **b)** Log2 FC from liver to colon epithelial cells in our scRNA-sea and the external scRNA-seq data plotted against each other. Colors as in a) with non-specific genes being depicted in gray. **c)** Log2 fold changes (FC) from metastasis to primary tumor samples of external microarray data stratified by the gene classification in Figure 5c. Fold changes of genes that are not differentially expressed in the microarray dataset (adjusted p-value < 0.05) are shown in black, other colors as in a). **d)** Log2 FC from metastasis to primary tumor samples of external microarray data of all genes that have been found to have a tissue-exclusive expression in our data. Colors as in a).

We further validated our findings using gene expression data from microarray sequencing of primary colon carcinomas and liver metastases from a cohort of 334 patients (184 primary colon carcinomas, 150 liver metastasis)^46–50^. After preprocessing and quality control of these data (Extended Data Figure 4b), we performed differential expression analysis between the primary tumor and metastasis samples (Methods, Extended Data Figure 4c). Genes with a differential expression in our scRNA-seq and a significant differential expression in the external microarray data showed highly consistent directions of effects for both the PT and MT candidate genes (Extended Data Figure 4d; 607 out of 909 genes measured in both data having the same direction of effect), as well as the PTCE and MTLE genes (Figure 4c; 321 out of 430 genes measured in both data having the same direction of effect). We could also confirm this agreement between our scRNA data and the microarray data for the tissue exclusive genes we defined (Figure 4d, Extended Data Figure 4e).

In conclusion, this analysis using two independent bulk- and single-cell datasets confirmed the adaptation of gene expression to the host tissue environment.

### Higher expressed pathways in tumor cells reflect surrounding expression signals

One example of a highly tissue-adaptive gene, which could be validated in the external data sets, is the small nuclear protein Prostate Cancer Susceptibility Candidate 1 (PRAC1). In our scRNA-seq data PRAC1 was detected in 44% of the primary colon cells and 63% of the colon epithelial cells, while it was not detected in metastasis and liver epithelial cells. This signal is also found in the external microarray data (log2FC MT/PT: −0.378, adj. p-value: 0.020). PRAC1 is specifically expressed in prostate, rectum, and distal colon^51^ and its expression is highly predictive of the side of the colon on which the cancer is located (more highly expressed in left-sided colon carcinomas)^52,53^. Thus, PRAC1 expression is highly tissue- and context-dependent, matching our findings of it being a primary tumor and colon epithelial exclusive gene, especially as our primary tumor samples were located on the left side of the colon. A tissue adaptive gene that was higher expressed in the liver epithelial and metastasis in both our dataset and the external data sets is the ERBB Receptor Feedback Inhibitor 1 (ERRFI1/Mig-6) (Our scRNA-seq data: log2FC MT/PT: 0.516, log2FC LE/CE: 1.463; external microarray data: log2FC MT/PT: 0.733, adj. p-value: < 0.001). ERRFI1 displays a strong tissue specific expression, being highly expressed in the liver and pancreas^54^, and is involved in the lipid metabolism, bile acid, and cholesterol synthesis in the liver^55^. An adaptive expression pattern in the metastasis may be crucial for the tumor cells to adapt their metabolism to the new environment.

To identify specific cellular functions affected by the tissue adaptation of gene expression we performed an analysis of pathways (BioCarta, KEGG, PID and Reactome, see Methods). Using Fast Gene Set Enrichment Analysis (FGSEA) we determined qualitative effect directions for each pathway and donor; that is, we determined if the pathway was more highly expressed in MT versus PT and - separately - if it was more highly expressed in LE versus CE. Subsequently, we defined pathways as ‘consistent’ if the effect directions of both MT-versus-PT and LE-versus-CE were the same across all donors. Of all pathways included in our analysis (n = 1103), 436 were ‘consistent’ according to this definition (Extended Data Table 6) Those 436 pathways showed a strong signal towards being upregulated either simultaneously in the primary tumor and colon epithelium (n = 234) or in the metastasis and liver epithelium (n = 147). Only very few pathways showed a consistent up- or down-regulation in the cross tissue pairs (higher in primary tumor and liver epithelial cells: 33, higher in metastasis and colon epithelial cells: 22). These findings are in line with our previous analysis (Figure 3a, 3c) and show that differences in expression patterns observed between the primary tumor and metastasis are to a large extent also observed in the epithelial cells and thus reflect cellular pathways that are differentially regulated depending on the tissue context.

Pathways reflecting tissue adaptive processes were clustered and annotated, resulting in five pathway groups higher expressed in metastases and liver epithelium and eight pathway groups displaying a higher expression in primary tumors and colon epithelial cells (Figure 5a). Biological processes linked to epithelial growth and cellular adhesion as well as extracellular matrix, cell-cell interaction and vesicular transport are observed to be higher expressed in the metastases and epithelial liver tissue. This makes sense in the context of the metastatic establishment in this novel tissue context and potentially necessary adaptation in the intercellular communication and transport. Other pathways displaying a higher expression in metastasis and liver are linked to different metabolic processes such as the lipid metabolism and oxidative stress response. Particularly noteworthy in this context is the pathway *Regulation of lipid metabolism by PPAR-α* that was consistently higher expressed in the metastasis and liver epithelial cells compared to the primary tumor and colon epithelium (Figure 5a). Importantly, these expression differences were validated using the external bulk- and single-cell expression data (Figure 5a, Extended Data Figure 5a). PPAR-α is a major regulator of the lipid metabolism in the liver, specifically regulating the fatty acid oxidation^43,44^. Further, the activation and expression of PPAR-α has been associated with tumorigenesis in colorectal cancer, as well as worse outcome in liver metastasis of colorectal cancer^56,57^. Thus, PPAR-α may be an important contributor for the adaptation of the tumor to the liver-specific lipid metabolism and establishing itself as a metastasis. Another pathway consistently higher expressed in the metastasis and liver epithelial in both the scRNA-seq and microarray data is the *Nrf2-ARE* pathway, an intrinsic mechanism of defense against oxidative stress^58^ (Figure 5b, Extended Data Figure 5b). The nuclear factor erythroid 2-related factor 2 (Nrf2) has been identified as both a tumor suppressor and an oncogene, as it can protect bening cells from oxidative stress^59^ as well as help cancer cells escape from reactive oxygen species (ROS)^60^.

**Figure 5:**
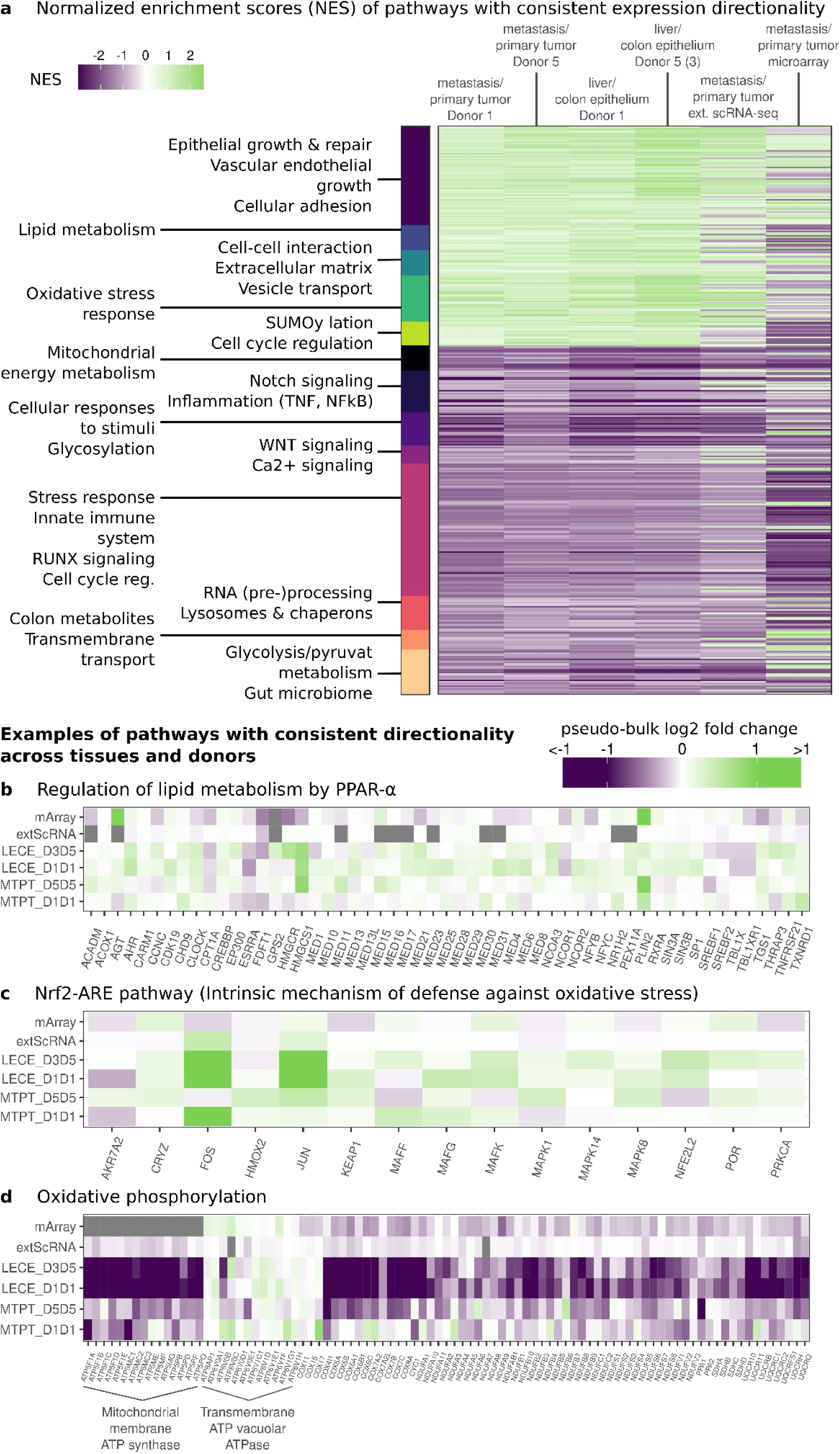
Pathways upregulated in PT or MT and the respective surrounding tissue. **a)** Normalized enrichment scores (NES) of all pathways with consistent directionality between MT & PT and LE & CE in our scRNA-seq data, clustered based on genes intersecting between different pathways. NES of pathway enrichment of external validation data are added. **b)** Log2 fold changes in between the primary tumor and metastasis cells (MTPT) as well as the liver & colon epithelial cells (LECE) of genes from the KEGG pathway ‘Regulation of lipid metabolism by PPAR-α’ which is higher expressed in the primary tumor and colon epithelial cells of our scRNA-seq data. Log2 fold changes in our scRNA-seq data, the external scRNA-seq data and external microarray data are displayed. All genes measured in our scRNA-seq data are visualized. **c)** As in b) but for the REACTOME pathway ‘Nrf2-ARE pathway’, which is higher expressed in the metastasis and liver epithelial cells of our scRNA-seq data. **d)** As in b) but for the KEGG pathway ‘Oxidative phosphorylation, which is higher expressed in the primary tumor and colon epithelial cells of our scRNA-seq data.

Two of the pathway groups higher expressed in the primary tumors and colon epithelium are closely linked to the function of the colon such as colon metabolite and transmembrane transport-related pathways as well as pathways connected to glycolysis/pyruvate metabolism and gut microbiome. These pathways are seemingly downregulated in the liver metastasis. Our previous finding, that genes related to the mitochondrial energy metabolism are upregulated in the primary tumor compared to the metastasis (Figure 2c, 2d) can also be observed in the colon epithelial compared to the liver epithelial cells, pinpointing to a context-driven energy metabolism of cancer cells. We could validate these results in independent scRNA-seq and microarray data (Figure 5d, Extended Data Figure 5c). These findings suggest that the shift in energy metabolism from oxidative phosphorylation to glycolysis commonly observed in liver metastasis may result from a metabolic adaptation to the new environment. Interestingly, while all detected subunits of the mitochondrial membrane ATP synthase (F1F0 ATP Synthase) were higher expressed in the cells derived from the colon site, the opposite was true for the vast majority of subunits of the transmembrane ATP vacuolar ATPase (V-ATPase). The F1F0 ATP Synthase generates ATP using a proton gradient and hence an increase in its subunits reflects an increase in energy availability through the mitochondria^61^. This may be a cellular adaptation to environmental changes such as alterations in nutrient availability, oxygen levels, or other external stress factors^62,63^. V-ATPases on the other hand acidify intracellular compartments by transporting proteins across the plasma membrane using ATP and play a fundamental role in maintaining pH homeostasis^64,65^. Therefore, they are involved in various physiological and pathological processes, such as macropinocytosis, autophagy, cell invasion, and cell death^65^. In this context, overexpression of a V-ATPase subunit and its regulators have been identified to being closely related to tumor invasion and metastasis, offering an explanation why we find the subunits of the V-ATPase being higher expressed in the liver epithelial and metastasis cells^65,66^.

## Discussion

The formation of metastases is the main cause of cancer-related death. The manifestation of metastases of a solid tumor requires cancer cells to change their characteristics and behavior, migrate out of the primary tumor, intra- and extravasate blood vessels and eventually proliferate in a different tissue^1–3^. Importantly, such adaptation may not require specific mutations to rewire metabolic networks. Instead, external cues, such as the availability of nutrients, oxygen and signaling molecules may suffice to adapt their metabolism^26,67,68^.

Here, we present a novel analysis strategy to identify tissue-adaptive gene expression changes that is generally applicable to metastases established from solid tumors. We reveal a pattern of tissue-adaptive expression changes, by utilizing a new scRNA-seq dataset with untreated, paired samples from cancer and benign tissue from donors with colorectal cancer and liver metastasis.

The upregulation of liver-specific pathways in liver metastasis indicates a metabolic adaptation having to occur in the metastatic cells. Our results strengthen the notion that epithelialization is required for the adaptation of colorectal metastasis to novel tissue environments^28^. We also find colon-specific pathways linked to e.g. the gut microbiome being highly expressed in the primary colon tumor but not in the liver metastasis. Taken together this indicates an adaptation of the expression pattern driven by external cues, which is consistent with the notion that the establishment of distant metastases from a primary colon tumor may not require *de novo* mutations^68,69^. Commonly reported differences in the expression pattern between the primary tumor and metastasis of specific cancer types, such as the switch from oxidative phosphorylation towards glycolysis in liver metastasis compared to primary colon tumor^26,38,39^, may be driven by tissue adaptive processes in the metastasis. The fact that only a small fraction of circulating tumor cells actually establishes a metastasis, makes it likely that the metabolic adaptation results from an interplay of genetic and epigenetic predisposition with signaling responses to external cues^67^.

Our work shows that such adaptation to a new host environment is not only required for a successful integration of the cancer cells into a novel tissue but may also contribute to the pathogenicity of a tumor. For example the upregulation of the *Nrf2-ARE* pathway that we have observed here can result from generally higher ROS levels in the liver compared to the colon, which is known to further increase the pathogenicity and treatment resistance of the metastasis^70–72^. Similarly, the activation and expression of PPAR-α, which is also classified as an tissue-adaptive process in our study, is associated with tumorigenesis in colorectal cancer, as well as worse outcome in liver metastasis of colorectal cancer^56,57^, resulting in PPAR-α being a repeatedly discussed cancer drug candidate^73,74^. Hence, targeting tissue-adaptive processes may not only prevent metastasization, but also address treatment resistances.

The general applicability of the approach presented here provides an opportunity for generating much deeper insight into the etiology of metastasis formation and the mechanisms determining treatment response way beyond colorectal cancer.

## Author contributions

LN, AB and AMH conceived this project. PSP, RW, MB, and CJB collected human tissue samples. AQ evaluated tissue histology. SH performed sample preprocessing, and flow sorting. MF performed scRNA sequencing. MW and LN performed quality control and filtering of the scRNA-seq data set. MW performed UMAP projection and cell type annotation. LN, MW, SH, MK, AMH and AB interpreted biological context. LN conceived and performed the DE analysis, the definition of tissue-adaptive and tissue-exclusive genes and interpretation of the results. LN performed FGSEA pathway analysis and interpretation. LN performed processing and analysis of the external scRNA-seq and microarray data sets. LN, SH, AB and AMH wrote the initial draft of the manuscript. All authors contributed to the writing of the final manuscript.

## Data availability

ScRNA-seq data counts and preprocessed count matrices will be made available over GEO.

## Code availability

Code for all analysis performed and the corresponding figures is provided on github: https://github.com/beyergroup/Gene-expression-adaptation-of-metastases.

## Human subject statement

The study was approved by the institutional review board of the University of Cologne (21-1467 and 21-1382_2). Informed consent was obtained from all subjects. Experiments conformed to the principles set out in the Declaration of Helsinki and the Department of Health and Human Services Belmont Report.

## Acknowledgement

We thank David Pertzborn, Anna Mühlig and Daniela Pelzel of the University Hospital Jena for discussion and support on validation work.

## Methods

### Sample collection and processing

Primary colon tumor (PT), normal colon (NC), liver metastasis (MT) and normal liver (NL) samples were collected from five donors with colorectal primary tumors and liver metastasis (Extended Data Table 1). All primary tumors were located at the colon sigmoideum or near rectum, except for donor 01 (D01), which had two primary tumors, one located in the sigmoideum and one in the ascending colon. The following samples were collected from each donor: D01: two PT, NC, MT, NL; D02: PT, NC; D03: MT, NL; D04: PT, NC, MT, NL; D5: PT, NC, MT, NL. Preparations of colon ascendence (PT) of D01 and MT of D04 resulted in too few cells for reliable analysis and were excluded after quality control from the study.

Fresh tissue samples from surgical resectates or endoscopies were dissociated as described earlier^75^. In brief, the tissue was minced to small pieces, disrupted in a C-tube used with the GentleMACS Dissociator (Miletnyi Biotec) combined with enzymatic digestion with DNAse I (500 U·mL−1; AppliChem PanReac), collagenase IV (320 U·mL−1; Thermo Fisher Scientific), and dispase II (2 U·mL−1; Sigma-Aldrich), filtered through a 100-µm cell strainer, collected and resuspended in 60% RPMI-1640 medium (Thermo Fisher Scientific), 30% FBS (Capricorn Scientific), and 10% dimethyl sulfoxide (DMSO) (Sigma-Aldrich) for freezing at −80°C. After 24 hours samples were transferred to liquid nitrogen until fluorescence activated cell sorting (FACS).

### FACS and single-cell sequencing

Single-cell suspensions were stained with PE/Cy7-conjugated antibodies against CD45 (Biolegend) as a leukocyte marker to reduce leukocytes and with propidium iodide (Thermo Fisher Scientific) to distinguish live and dead cells according to the manufacturer’s specifications. In corporation with the FACS & Imaging Core Facility of the Max Planck Institute for Biology of Ageing, Cologne, flow cytometry assisted cell sorting was performed on BD FACSAria Fusion (BD Biosciences) using a 100-µm nozzle. For each sample, 10.000 living cells were sorted, comprising 30% of CD45-positive cells and 70% of CD45-negative cells, whenever applicable. For samples of D04 and D05, an additional staining for EpCAM using CD326 (EpCAM) FITC Antibody anti-human, clone HEA-125 (Miltenyi Biotec), was performed to further enrich for epithelial cells aiming at a ratio of ⅓ CD45+, ⅓ EpCAM+, and ⅓ CD45-EpCAM-cells. Collected single cells were placed on ice and further processed using the Chromium™ Single Cell 3’ Solution (10x Genomics) and sequenced on a NovaSeq 6000 (Illumina) with Illumina 3’ v3.1-paired end chemistry, 29-10-10-89 bp aiming at 25,000 read pairs per cell. Targeted cell recovery was aiming at 3000 cells per sample.

### Raw data analysis and quality control

Raw single cell data were first analyzed with the *Cell Ranger* pipeline from 10x Genomics^76^. The filtered feature matrices were read in to analyze the data using Seurat (version 4.0.3)^77–80^, excluding genes that were expressed in less than three cells and excluding cells that expressed less than 1000 different genes. The quality was assessed for each sample separately, excluding cells that had a higher number of detected genes or UMIs than a respective threshold, defined by either three median absolute deviations from the median or three standard deviations from the mean, choosing the more stringent of the two for every individual cell. Initially, all cells with a fraction of mitochondrial genes (*percMito*) higher than 30% regarding total UMIs were excluded. After preliminary clustering the top 15% of cells regarding the percMito were additionally excluded due to a cell type dependency, allowing an individual cell type specific threshold for this parameter, thus taking physiological differences between cell types into account^81^.

### UMAP projection and cell type annotation of scRNA-seq

The filtered data from above was further normalized for UMAP projection, clustering and cell type annotation. Samples that were sequenced in the same run were merged, normalize (*Seurat::NormalizeData* with *normalization.method=‘LogNormalize’* and scaled (e.g. the overall expression of a gene among all cells was transformed to a mean expression of 0 with a variance of 1), according to the workflow provided by the Satija Lab^82^. The samples were integrated for the UMAP projection using Canonical Correlation Analysis to match similar cell types from different batches (built-in Seurat method)^77^. The most variable features were identified(*Seurat::FindVariableFeatures* with *selection.method=’vst’, nfeatures=6000, assay=‘RNA’*) Then principal component analysis (PCA) was performed. Cells were clustered following the analysis pipeline of the developers of Seurat^79^ and Luecken & Theis^83^, using the clustering algorithm of the Seurat platform, including a MMN-graph and Louvain algorithm. For that, a K-nearest neighbor graph based on Euclidean distance in the space defined by PCA and the number of principal components was constructed, with edge weights based on Jaccard similarity (i.e. being neighbors in space)^84^. The Louvain algorithm was then used to iteratively group cells together with the goal of optimizing the standard modularity function^85^. The Uniform Manifold Approximation and Projection (UMAP) algorithm^86,87^ was used to generate a non-linear visualization of the data.

Differential expression analysis to find top marker genes for each cluster was performed using the previously defined most variable features, the algorithm MAST^88^, a minimum fraction of cells expressing the differentially expressed genes of 0.4 and a minimum log-fold change in expression of 0.1. Cell types were defined using common surface markers and top differentially expressed genes per cluster with the help of PanglaoDB^89^, a public database of a variety of scRNA-seq experiments which allows to browse gene expression and cell type annotations from published data sets. EPCAM was used to identify epithelial cells. Among EPCAM-positive cells, cells of cancer origin were discriminated using inferCNV^90^ (settings as recommended by the developers). For immune cells, CD79A was used to identify B cell species and NKG7, TRBC2 and IL7R for T cells. PDGFRA, collagen species and matrix-specific genes discriminated against fibroblast species. VWF was used as a marker of endothelial cells.

### Defining differentially expressed genes between cell type groups

We developed a pipeline to estimate differential expression changes, which delivers stable and unbiased results even if there are big differences in the sequencing depth between the groups of cells we are aiming to compare (Extended Data Figure 2a).

Starting with the filtered but not normalized or integrated expression values, cells were divided into cell groups based on their cell type and donor they originate from. Cell groups containing less than 49 cells were removed from the analysis. Next, all genes which were not expressed in at least 10% of the cells per cell group contained in the analysis are removed (6872 genes remained in the analysis). Since genes with many zeros were removed, we assumed that the vast majority of the remaining genes are actually expressed and hence only consider gene expression >0 into account, allowing us to better account for big differences in the sequencing depth. Therefore, after log2 transforming the data, the mean expression of every cell group was estimated, without taking the zeros into account. Subsequently, the mean expressions per cell group were centered, setting the library size of every cell group to 10,000. Log2 fold changes (log2 FC) were estimated donor-wise, using the mean expressions per cell group, always pairing the donor matching cell groups to each other, resulting in donor-specific normalized log2 FC for donor 1 and donor 5. Genes with a normalized log2 FC greater than 0.1 or lower than −0.1, which agree in directionality of the log2 FC in all donor-specific groups are defined as candidate genes. 530/6872 genes were defined to be primary tumor specific, while 305/6872 were metastasis specific.

The approach was additionally utilized to compare not only metastasis and primary tumor but also liver epithelial cells with colon epithelial cells. Cell groups of all four cell types were defined for donor 1 and donor 5, except for the cell group of liver epithelial cells of donor 5, since there were none sequenced. This group was subsidized with epithelial liver cells of donor 3. 5019 genes remained in the analysis after removing genes which were too lowly expressed (<10%) in at least one of the cell groups. Since the gene filtering in the bioinformatic pipeline is influenced by tissues used in the individual analysis, the genes used for defining PT and MT candidates (Figure 2a, b) slightly differ from those included in PTCE and MTLE analysis (Figure 3b, c). This influences the pseudo-bulk expressions and therefore result in minimal different log2 fold changes estimated between metastasis and primary tumor cells (Extended Data Figure 4). Due to those differences the PTCE and MTLE candidates are not just a subset of the PT and MT candidates even though there is a high overlap.

### Defining tissue exclusively expressed genes between cell groups

When defining differentially expressed genes, genes which show an expression below 10% in any of the defined cell groups are excluded, leading to the analysis overseeing potentially very interesting genes; genes which are not at all or almost not at all expressed in one cell group but highly expressed in the one we want to compare it to. Therefore, we additionally defined exclusively expressed genes based on the same cell groups used when inferring differentially expressed genes. For each gene the percentage of times it was expressed in each cell group was estimated and subsequently the weighted mean over the cell groups from the same cell types was retrieved. Genes expressed in under 10% of the genes in one cell type and over 20% in the other cell type are defined to be expressed exclusively.

### FGSEA pathway enrichment on cell type groups

Pathway enrichment was performed utilizing fast gene set enrichment analysis (FGSEA) in the form of the R package *fgsea* (version 1.31.0)^91^, setting the minimum node size to 10. Canonical pathways^92^ (accessed on 03.01.2023) from BioCarta, KEGG, PID and Reactome were provided to the algorithm. FGSEA was applied on the log2 normalized mean expressions estimated for each donor included in the analysis. Pathways were defined as primary tumor & colon epithelial or liver metastasis & liver epithelial specific based on identical directionality of the enrichment between the donors. These specific pathways were clustered by using k-means^93^ after estimating a distance matrix based on the number of genes pathways share (Jaccard index)^94^ and performing Kruskal’s Non-metric Multidimensional Scaling^95^ (*MASS:isoMDS*, package version 7.3-61)^96^.

### Differential expression analysis in external scRNA-seq data

Public available external scRNA-seq data from matching primary colon tumor and liver metastasis was downloaded (n = 10, preprocessing and cell type annotation as provided by the authors, GEO accession: GSM7058755)^45^. Differential expression analysis was performed with the same pipeline implemented for our scRNA-seq data. Cells were divided into cell groups based on their cell type and donor they originate from and containing less than 120 cells were removed from the analysis. Next, all genes which were not expressed in at least 5% of the cells per cell group contained in the analysis are removed (6652 genes remained in the analysis). The data was log2 transformed and mean expression of every cell group was estimated (without taking the zeros into account) and centered (setting the library size of every cell group to 10,000). Mean pseudo-bulk metastases and primary tumor expression values were estimated and subsequently the overall mean log2 fold change (MT/PT) was estimated. Pathway enrichment was performed on the log2 fold changes by applying FGSEA, using the same set of pathways we enriched in our scRNA-seq data.

### Differential expression analysis in external microarray data

Microarray data from two primary colon tumor data sets and one liver metastasis data set was preprocessed together (GEO accession: GSE79959, GSE96528, GSE79959)^46–50^. Datasets with over 150 samples were subsampled to a maximum of 150 samples per dataset (Extended Data Table 7) and subsequently preprocessed together via robust multi-average normalization using the *oligo* R package (version 1.68.1)^97^. Principal component analysis (PCA) revealed an overlap of the primary tumor samples in the first and second PC while the samples of the liver metastasis separated from them (Extended Data Figure 2a). Differential expression analysis between the metastasis and primary tumor was performed using the pipeline implemented by *limma* (version: 3.58.1)^98^, resulting in estimated log2 fold changes and adjusted p-values. Pathway enrichment was performed on the log2 fold changes by applying FGSEA, using the same set of pathways we enriched in our scRNA-seq data.

## Extended Data Figures

**Extended Data Figure 1:**
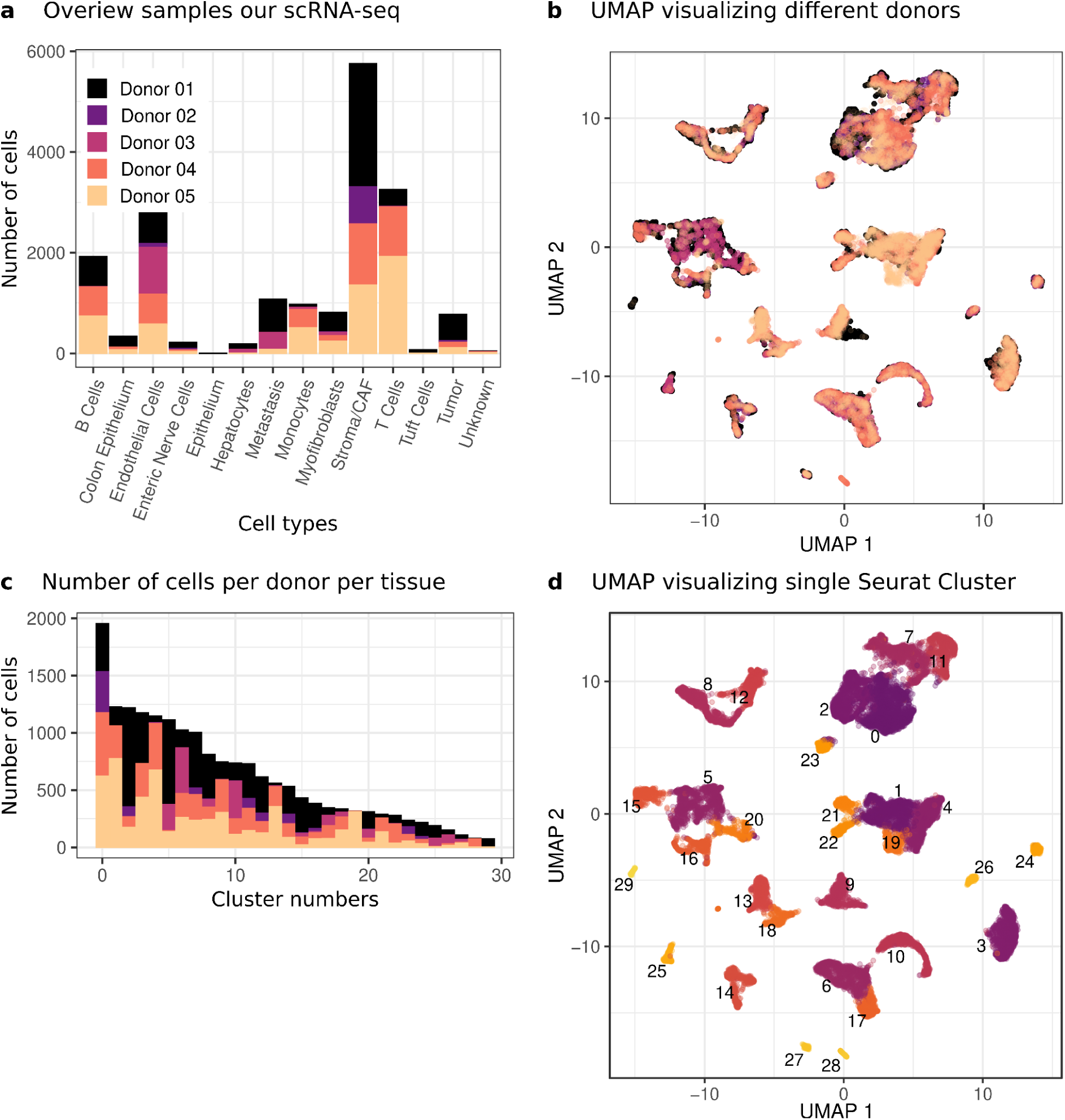
Overview of scRNA-seq dataset from five donors with primary colon tumor and liver metastasis. **a)** Barplot showing the number of cells per cell type colored by donor. **b)** UMAP of scRNA-seq data with colors indicating different donor origin/sample site origin. **c)** Barplot showing the number of cells per cell type per donor. **d)** UMAP of scRNA-seq data with colors indicating different cell types.

**Extended Data Figure 2:**
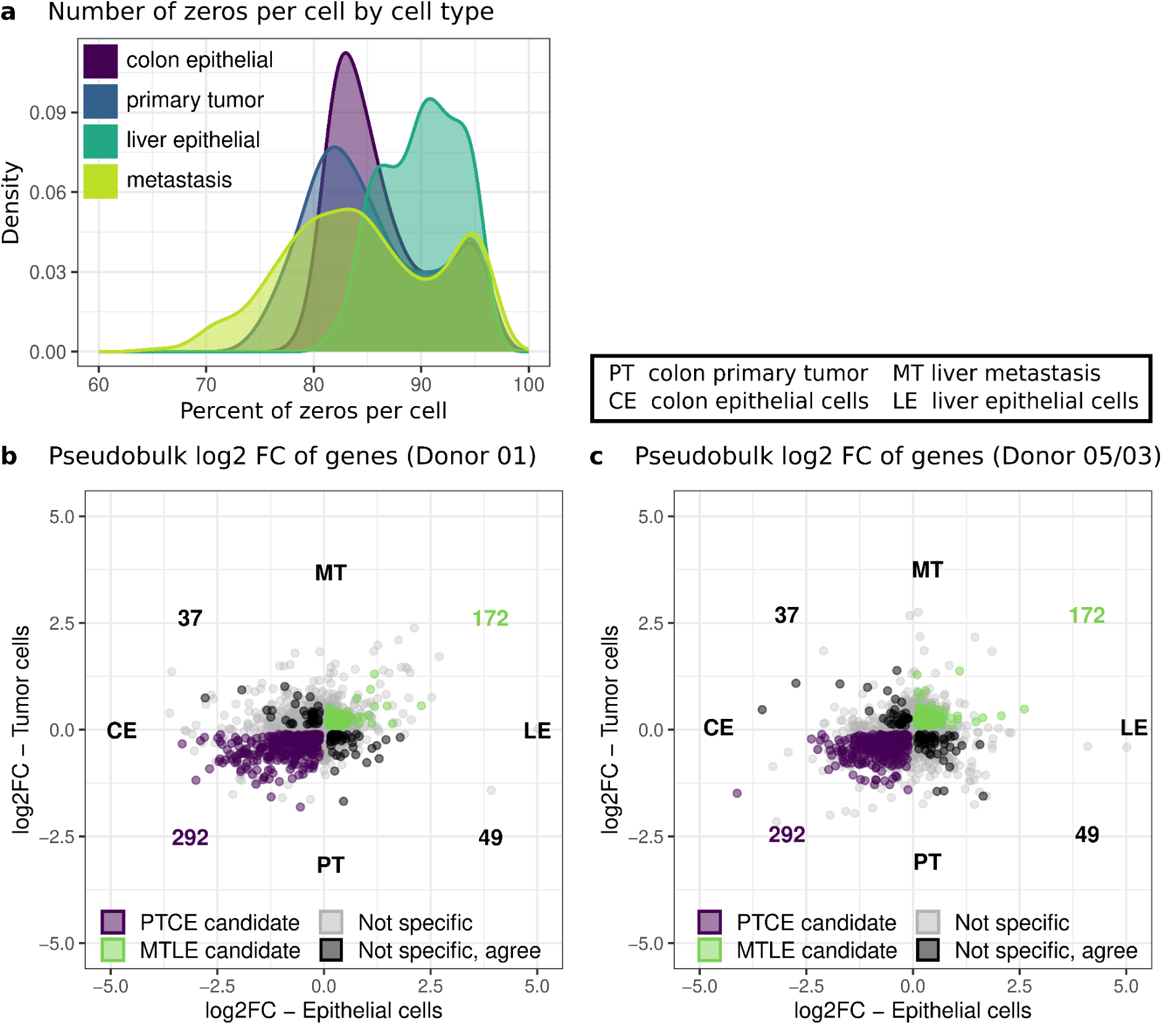
Differences in expression between MT and PT as well as LE and CE cells. **a)** Percent of zeros per cell (after filter for low quality cells, see methods) for colon epithelial (purple), colon primary tumor (blue), liver epithelial (turquoise) and liver metastatic (light green) cells over all donors. **b)** Mean log2 fold change of metastasis and primary tumor plotted against the mean log2 fold change of liver and colon epithelial cells of donor 1. Genes with expression patterns consistent between the donors are highlighted (primary tumor and colon epithelial specific genes = purple, metastasis and liver epithelial specific genes = green, other = black, four outlier genes are not visualized in the plot). **c)** As b) but for donor 5 (liver epithelial cells from donor 3).

**Extended Data Figure 3:**
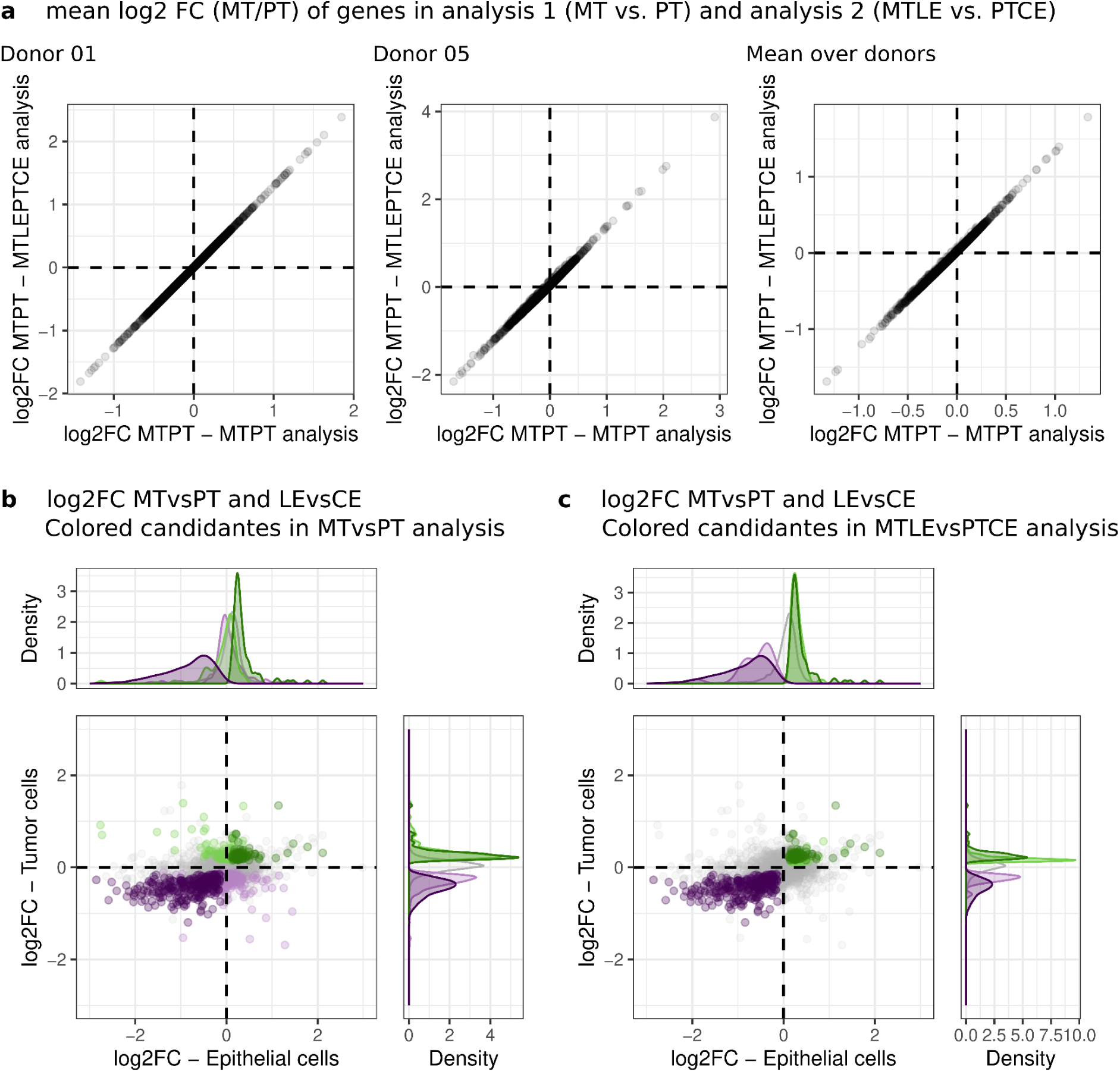
Differences in analysis and candidates between MT vs PT and MTLE vs PTCE analysis. **a)** Log2 FC of primary tumor vs metastasis comparison (Figure 2) and primary tumor vs metastasis & liver vs colon epithelial cells comparison (Figure 3) plotted against each other in donor 1 (left), donor 5 (middle) and mean over all donors (right). **b)** Mean log2 fold change of metastasis vs primary tumor plotted against the mean log2 fold change of liver vs colon epithelial cells estimated over the donors. Genes, which are defined as both, cancer specific (Figure 2b) and cancer & surrounding tissue specific (Figure 3c), are depicted in dark purple (primary tumor & primary tumor + colon epithelial specific) and dark green (metastasis & metastasis + liver epithelial specific), while genes which are only cancer specific defined by the metastasis vs primary tumor analysis (Figure 2b) are light purple (primary tumor) and light green (metastasis). Genes with no specificity are gray. (Four outlier genes are not visualized in the plot.) **c)** As b) but light colored genes are genes which are only cancer & surrounding tissue specific (Figure 3c, light purple = primary tumor & colon epithelial, light green = metastasis & liver epithelial).

**Extended Data Figure 4:**
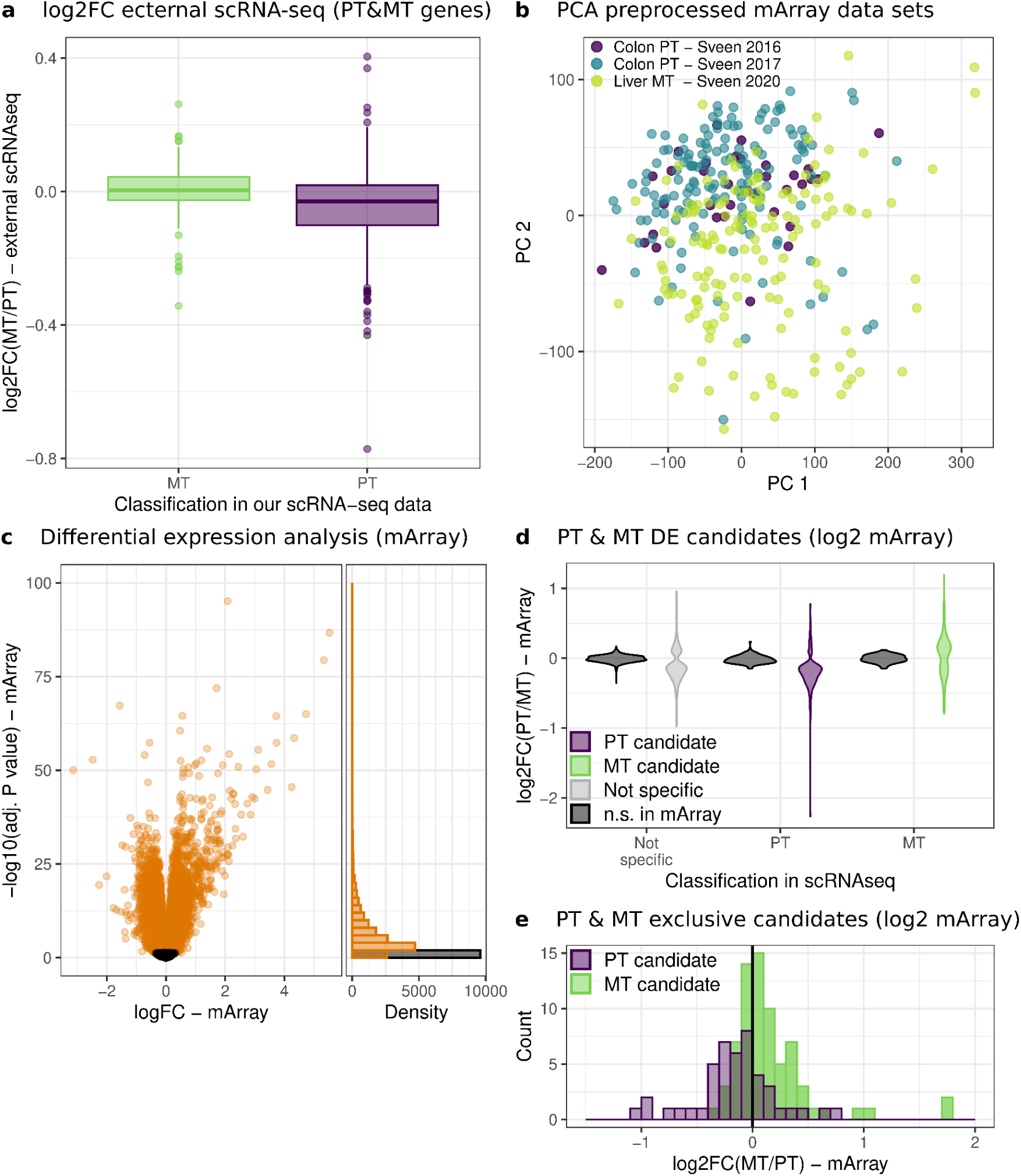
Validation of tissue specific candidate genes with external data sets. **a)** Log2 FC from metastasis to primary tumor cells in external scRNA-seq data plotted by their classification in our data (PTCE candidates = purple, MTLE candidates = green). **b)** PCA of preprocessed microarray data sets, colored by dataset and sample origin. **c)** Log2 FC and adjusted p-values of external microarray data from MT compared to PT. Genes that are differentially expressed (adjusted p-value =< 0.05) are depicted in orange, all others in black. **d)** Log2 FC from MT to PT samples of external microarray data set plotted by their classification in our data (PT specific = purple, MT specific = green, non-specific = gray). Genes which are not differentially expressed in the microarray dataset (adjusted p-value < 0.05) are colored in black. **e)** Log2 FC from MT to PT samples of external microarray data set of all genes which have been found to have a tissue exclusive expression in our data (PT exclusive = purple, MT exclusive = green).

**Extended Data Figure 5:**
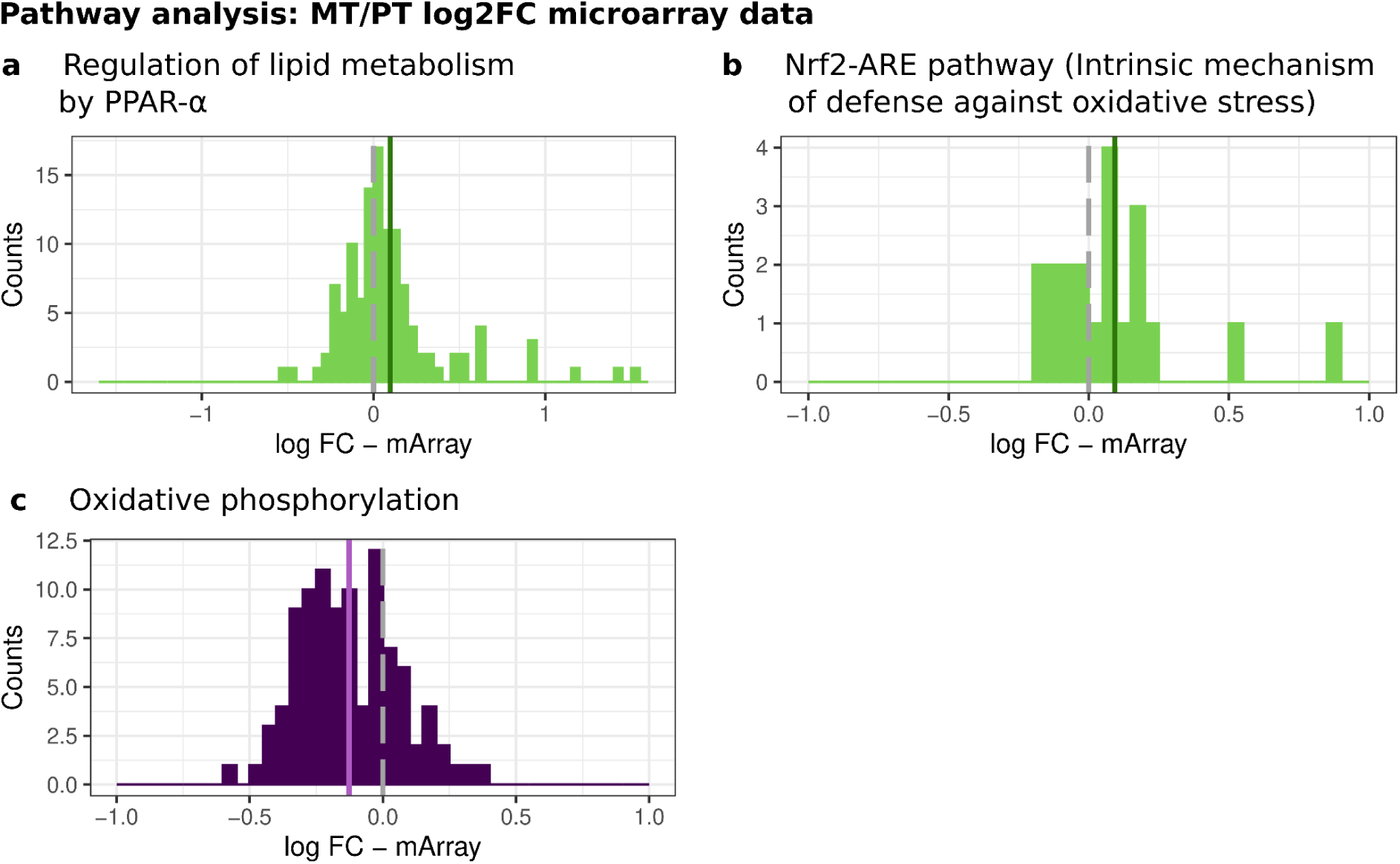
Pathway analysis. **a)** Log2 fold changes in between the primary tumor and metastasis cells (MT/PT) in external microarray data for all genes involved in the KEGG pathway ‘Regulation of lipid metabolism by PPAR-α’, which is higher expressed in the metastasis and liver epithelial cells of our scRNA-seq data. The mean gene expression of all pathway genes in the external microarray data is visualized with a dark green line. **b)** As in a) but for the REACTOME pathway ‘Nrf2-ARE pathway’, which is higher expressed in the metastasis and liver epithelial cells of our scRNA-seq data. The mean gene expression of all pathway genes in the external microarray data is visualized with a dark green line. **d)** As in a) but for the KEGG pathway ‘Oxidative phosphorylation’, which is higher expressed in the primary tumor and colon epithelial cells of our scRNA-seq data. The mean gene expression of all pathway genes in the external microarray data is visualized with a light purple line.

## Extended Data Tables

**Extended Data Table 1: Human donor and sample information.**

**Extended Data Table 2: Overview over all cells after quality control.** Quality criteria, sample and cell information as well as UMAP projection are provided.

**Extended Data Table 3: DE comparison of liver metastasis vs. primary colon tumor cells.** All genes included in the analysis are provided. Log2 FC of the individual donors a the mean over all donors and a binary classification if a gene is metastasis or primary tumor specific are provided for every gene. Log2 FC of the external scRNA-seq data and log2 FC and adjusted p-value of external microarray data analysis are included.

**Extended Data Table 4: DE comparison of liver metastasis vs. primary colon tumor and liver vs. colon epithelial cells.** All genes included in the analysis are provided. Log2 FC of the individual donors a the mean over all donors and a binary classification if a gene is metastasis vs primary tumor specific, liver vs. colon epithelial specific or specific for both are provided for every gene. Log2 FC of the external scRNA-seq data and log2 FC and adjusted p-value of external microarray data analysis are included.

**Extended Data Table 5: Overview over ‘tissue-exclusive’ genes.** Percent of cells in which gene is expressed per cell type is provided and a binary classification if a gene is metastasis vs primary tumor specific or metastasis vs. primary tumor AND liver vs. colon epithelial specific provided for every gene. Log2 FC and adjusted p-value of external microarray data analysis are included.

**Extended Data Table 6: Pathway enrichment liver metastasis vs. primary colon tumor and liver vs. colon epithelial cells.** All pathways included in the analysis are provided. Normalized enrichment scores and adjusted p-values of the individual donors as well as the mean over all donors and a binary classification if a gene is metastasis vs. primary tumor specific or metastasis vs. primary tumor AND liver vs. colon epithelial specific are provided for every pathway. Normalized enrichment scores and adjusted p-values of pathway enrichment of the external scRNA-seq data and external microarray data analysis are included.

**Extended Data Table 7: List of microarray samples used from the external microarray data sets** (see Methods).

